# Horizontal gene transfer of a unique *nif* island drives convergent evolution of free-living N_2_-fixing *Bradyrhizobium*

**DOI:** 10.1101/2021.02.03.429501

**Authors:** Jinjin Tao, Sishuo Wang, Tianhua Liao, Haiwei Luo

**Author notes:** These authors contribute equally to this work (Jinjin Tao, Sishuo Wang). **Corresponding author:** Haiwei Luo, School of Life Sciences, The Chinese University of Hong Kong, Shatin, Hong Kong SAR.

## Abstract

The alphaproteobacterial genus *Bradyrhizobium* has been best known as N_2_-fixing members that nodulate legumes, supported by the *nif* and *nod* gene clusters. Recent environmental surveys show that *Bradyrhizobium* represents one of the most abundant free-living bacterial lineages in the world’s soils. However, our understanding of *Bradyrhizobium* comes largely from symbiotic members, biasing the current knowledge of their ecology and evolution. Here, we report the genomes of 88 *Bradyrhizobium* strains derived from diverse soil samples, including both *nif*-carrying and non-*nif*-carrying free-living (*nod* free) members. Phylogenomic analyses of these and 252 publicly available *Bradyrhizobium* genomes indicate that nif-carrying free-living members independently evolved from symbiotic ancestors (carrying both nif and nod) multiple times. Intriguingly, the *nif* phylogeny shows that all *nif*-carrying free-living members comprise a cluster which branches off earlier than most symbiotic lineages. These results indicate that horizontal gene transfer (HGT) promotes *nif* expansion among the free-living *Bradyrhizobium* and that the free-living *nif* cluster represents a more ancestral version compared to that in symbiotic lineages. Further evidence for this rampant HGT is that the *nif* in free-living members consistently co-locate with several important genes involved in coping with oxygen tension which are missing from symbiotic members, and that while in free-living *Bradyrhizobium nif* and the co-locating genes show a highly conserved gene order, they each have distinct genomic context. Given the dominance of *Bradyrhizobium* in world’s soils, our findings have implications for global nitrogen cycles and agricultural research.

## Introduction

Members of the slow-growing genus *Bradyrhizobium* constitute an important group of rhizobia [1, 2]. They symbiotically fix N_2_ with diverse legume tribes and are the predominant symbionts of a wide range of nodulating legumes [3, 4]. Recent studies, however, have provided accumulating evidence for the existence of close relatives of rhizobia including *Bradyrhizobium* that do not carry *nod* genes which are essential for the establishment of nodulation, and thus are not able to nodulate legumes [5]. Increasingly, *Bradyrhizobium* strains without *nod* have been found to be particularly abundant in soils [6–8]. VanInsberghe et al [8] showed that non-symbiotic *Bradyrhizobium* was the dominant bacterial lineage in North America coniferous soils. A global atlas of the dominant soil-dwelling bacteria based upon 16S rRNA gene amplicon sequencing indicated that the genus *Bradyrhizobium* was the most abundant bacterial lineage in soils across the world [6]. A recent study suggested that ancestral lineages of *Bradyrhizobium* might adapt to a free-living lifestyle, highlighting the importance of the free-living lifestyle in the evolution of *Bradyrhizobium* [9].

Although the majority of N_2_ fixation in terrestrial ecosystems is generally thought to be performed by symbiotic rhizobia [10–12], free-living N_2_-fixing bacteria may be important contributors to the nitrogen budgets in a number of environments, for example soil ecosystems that lack leguminous plants [13]. Free-living N_2_ fixation also occurs in understudied ecosystems such as deep soil and canopy soil [14, 15], which may lead to the underestimation of its global rates. Rhizobia are generally thought to be capable of N2 fixation only in nodules [16, 17]. The exceptions are found in some members of *Bradyrhizobium* and *Azorhizobium* which were shown to be able to fix N_2_ in the free-living state [18, 19]. Such cases in *Bradyrhizobium* were mostly explored in photosynthetic members [18, 19], but have recently been found in other strains. For instance, *Bradyrhizobium* sp. AT1, a non-symbiotic strain isolated from the root of sweet potato was reported to fix N_2_ with a rate comparable to the photosynthetic strain *Bradyrhizobium* sp. ORS278 under the molecular oxygen (O_2_) concentrations at 1-5% (for both) in the free-living state [20]. Another example is *Bradyrhizobium* sp. DOA9, which harbors a *nif* cluster on its chromosome and another on a symbiotic plasmid: while both could participate in N_2_ fixation during symbiosis, only the chromosomal one participated in N_2_ fixation in the free-living state [21].

Compared with the nodules, the condition of soils is harsh for N_2_ fixation for several reasons. First, in contrast to being at nanomolar levels in nodules, the O_2_ concentration in soils could be up to atmospheric levels [22], which is highly detrimental to N2 fixation, a process that requires an anaerobic microenvironment [23]. Second, unlike symbiotic rhizobia, free-living rhizobia lack readily available organic matter provided by the host plants in exchange for fixed nitrogen. Last, free-living soil bacteria are frequently exposed to stresses like droughts, high osmolarity, and high temperature. It is therefore imperative to investigate the strategies that free-living N_2_-fixing *Bradyrhizobium* may use to deal with the harsh condition in soils.

Despite the potential ecological significance, free-living *Bradyrhizobium*, particularly those able to fix N_2_, have been poorly characterized. To this end, we isolated 88 *Bradyrhizobium* and five related strains from soils of soybean cropland, artificial park, forest, and grassland at several geographic locations in China, and had their genomes sequenced. By building a phylogenomic tree and reconstructing the ancestral lifestyles with additional 252 genomes where 42 are free-living *Bradyrhizobium* strains deposited in public databases, we revealed complex lifestyle shifts along the evolutionary history of *Bradyrhizobium*. Notably, we showed that horizontal gene transfer (HGT) of a unique *nif* island among free-living members may have facilitated their transitions from symbiotic lifestyle and adaptation to soil habitats.

## Methods and Materials

Detailed methods are described in Supplementary Text S1. In brief, we collected soil samples from five different sites in China (Fig. S1A) that cover several ecosystem types: soybean cropland (Heihe and Lvliang), artificial park (Shenzhen), undeveloped forest (Lanzhou), and grassland (Hefei) (see Fig. S1A and Data Set S1 for details). Five different media were applied to retrieve *Bradyrhizobium* from the samples. We obtained isolates that likely resembled *Bradyrhizobium*, and had their genomes sequenced with the Hiseq X platform (Illumina), followed by genome assembly and gene identification using SPAdes v3.10.1 [24] and Prokka v1.12 [25], respectively. Completeness of these genomes was calculated using CheckM v1.0.7 [26] with default parameters.

The phylogenomic tree of *Bradyrhizobium* was constructed using IQ-Tree v1.6.2 [27] with the 93 strains isolated in the current study and 252 genomes deposited in the NCBI Genbank database based on amino acid sequences of 123 shared single-copy genes identified by OrthoFinder v2.2.7 [28]. *Bradyrhizobium* members were divided into three lifestyle types based on the presence or absence of *nod* and *nif* genes (Data Sets S1 and S2). Strains that possess *nifBDEHKN* and *nodABCIJ* genes (at most one missing gene allowed; the same thereafter) were classified as symbiotic (Sym), strains possessing only *nifBDEHKN* genes were classified as free-living N_2_-fixing (FL_nif_), and those lacking both *nod* and *nif* genes were classified as free-living non-N_2_-fixing (FL_nonnif_). The lifestyle of ancestral nodes in the phylogenomic tree was inferred using Mesquite v3.61 [29] by the maximum parsimony reconstruction method, which infers the ancestral lifestyles by minimizing the number of steps of lifestyle change along the phylogenomic tree. Similar to a recent study [9], we did not apply the methodologically complex maximum likelihood method because the frequent lifestyle transitions across short branches in the *Bradyrhizobium* phylogeny likely leads to overestimates of the transition rate and hence inaccurate reconstruction of ancestral lifestyles.

Metadata of 4,958 sequence files (runs) were retrieved from the NCBI Sequence Read Archive (SRA) using the term “*nifH* metagenome” (last accessed in December 2020). Raw data were downloaded using the SRA Toolkit (https://github.com/ncbi/sra-tools) and were further processed with QIIME2 [30]. We applied evolutionary placement methods to assign the short sequence reads to the four phylogenetically distinguishable types of *nif* clusters (i.e., FL, PB, Sym and SymBasal; see Fig. 2) using PaPaRa v2.5 [31] and EPA-ng v0.3.8 [32]. The normalized abundance of each type of *Bradyrhizobium nifH* genes was calculated as the number of reads assigned to the corresponding *nifH* type divided by the total number of reads in the amplicon sequencing data set assigned to *Bradyrhizobium*.

## Results and discussion

### Newly sequenced free-living strains expand ecological diversity of *Bradyrhizobium*

We isolated and sequenced the genomes of 93 *Bradyrhizobium* strains initially identified by 16S rRNA gene (five were later shown to belong to *Afipia* based on phylogenomic construction; see below) isolated from diverse types of soils, i.e., lateritic red earths, black soils, brown coniferous forest soils, cinnamon soils, and yellow-brown earths in different sites of China (Figure S1). Among these isolates, 81 were non-symbiotic strains, as evidenced by the lack of the nodulation-determining *nod* genes, four of which carried *nif* genes. The remaining twelve were potential symbiotic strains that were likely released from decaying nodules: 11 of them carried *nod* genes and the other one phylogenetically clustered with photosynthetic members which use a nod-independent strategy for nodulation [33] (Fig. 1); all of them were isolated from the rhizosphere or bulk soil of leguminous plants (Data Set S1). Together with the 215 publicly available *Bradyrhizobium* genomes, our data set included 167 symbiotic (Sym), 22 free-living N_2_-fixing (FL_nif_), and 119 free-living non-N_2_-fixing (FL_nonnif_) strains (Data Sets S1 and S2).

**Figure 1.**
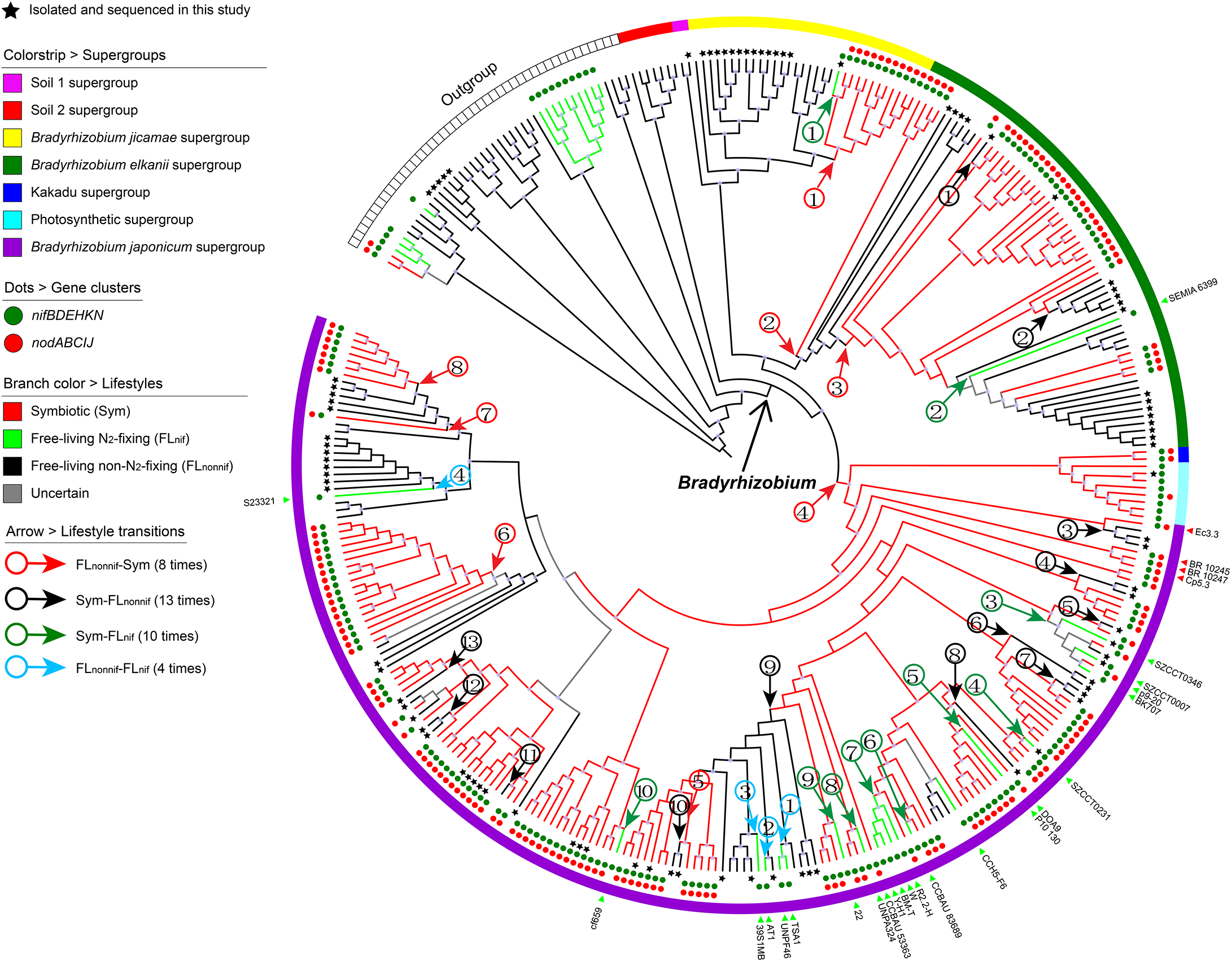
The maximum-likelihood phylogenomic tree of *Bradyrhizobium* and inferred lifestyle evolutionary history. Strains from *Xanthobacteraceae* were used as an outgroup. Ancestral lifestyles were inferred using the parsimony method in Mesquite. Branches in black, green, red and gray represent free-living non-N_2_-fixing, free-living N_2_-fixing, symbiotic and uncertain lifestyles, respectively. The color strips in the outer layer indicates different supergroups. Red and green solid circles denote the presence of *nod* and *nif* genes, respectively. Black stars adjacent to the tips of the phylogeny indicate strains isolated and sequenced in the present study. We inferred 8, 13, 10 and 4 times FL_nonnif_-Sym, Sym-FL_nonnif_, Sym-FL_nif_ and FL_nonnif_-FL_nif_ transitions respectively (noted with arrows). Purple circles on the nodes indicate ultrafast bootstrap values higher than or equal to 95% calculated by IQ-Tree. The red and green triangles on the outer layer indicate symbiotic members of the basal lineages of the *B. japonicum* supergroup and free-living N_2_-fixing strains, respectively.

**Figure 2.**
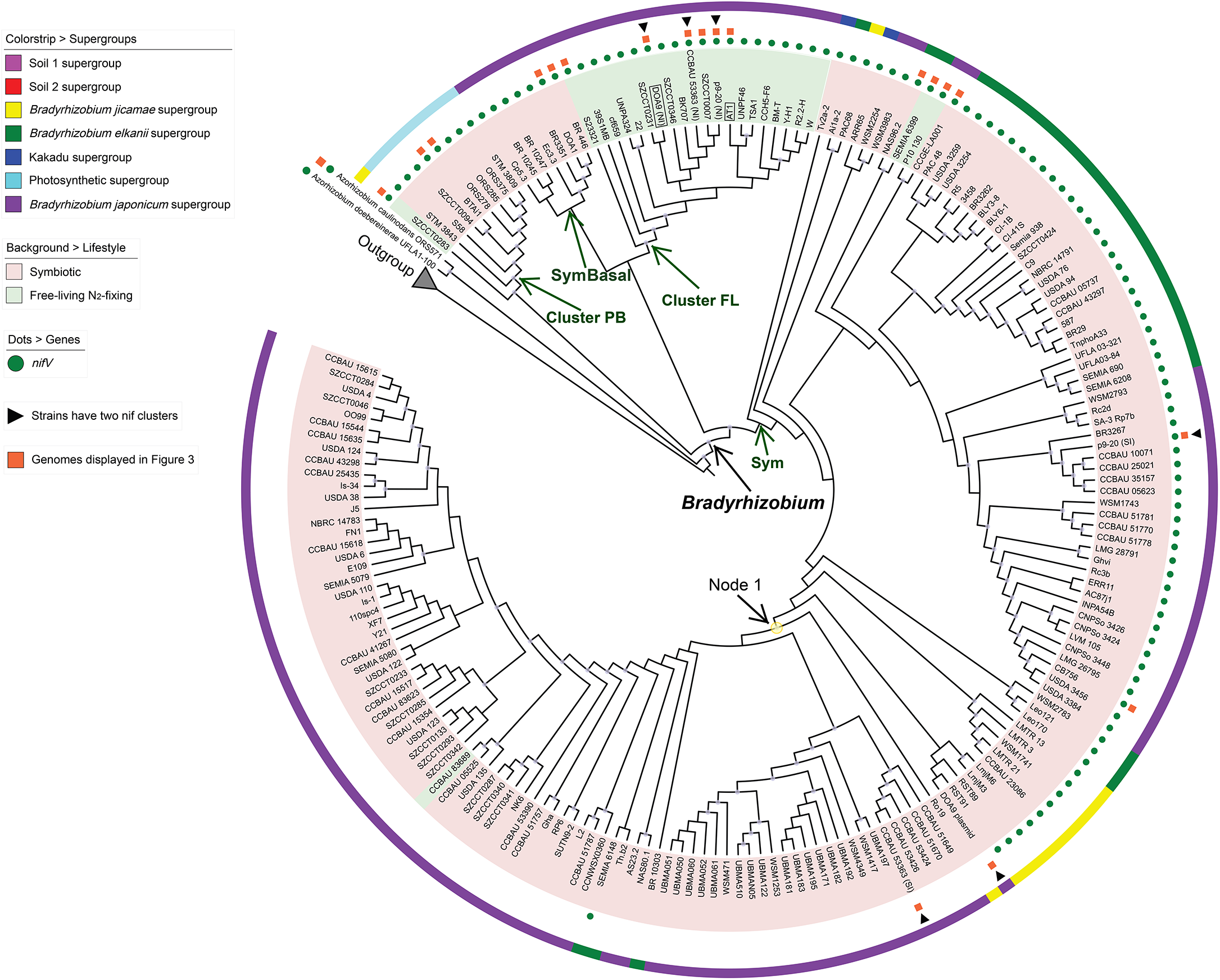
*Nif* gene phylogeny of *Bradyrhizobium*. The phylogeny was constructed using the concatenated alignments of *nifABDEHKNVX*. *Nif* genes from *Rhizobium*, *Sinorhizobium*, *Mesorhizobium*, *Azorhizobium* and *Beta*-rhizobia (rhizobia from *Betaproteobacteria*) are used as the outgroup. The color strips on the outer layer indicates different supergroups, and the background colors of the species name represent different lifestyles. The purple circles on the nodes indicate ultrafast bootstrap values higher than or equal to 95% calculated by IQ-Tree. The three symbiotic strains (also mentioned in Fig. 1) with *nif* genes on both the free-living *nif* island and symbiosis island are indicated by black triangles. The tips with a black box around denote the strains from Cluster FL with experimental evidence for their N_2_ fixation in the free-living state. Note that three symbiotic strains, namely DOA9, p9-20 and CCBAU 53363, carried a *nif* cluster similar to the one on the free-living *nif* island (NI) in addition to another one on the symbiosis island (SI). The geographic locations of free-living N2-fixing strains are displayed in Fig. S1B.

Despite a significantly expanded data set, all the *Bradyrhizobium* isolates fell into seven phylogenetic supergroups (Fig. 1) named in a recent study [2]. These were the Soil 1, Soil 2, *B, jicamae*, *B. elkanii*, Kakadu, Photosynthetic, and *B. japonicum* supergroups (Fig. 1). Among the 93 newly isolated strains, 88 contributed to all these supergroups except for the Soil 2 and Kakadu supergroups, and the remaining five strains were related to the genus *Afipia* in the outgroup. The newly isolated strains were phylogenetically nested within those that have publicly available genomes, except for the 13 strains in the *B. jicamae* supergroup which formed a sister clade to the previously sequenced strains of that supergroup (Fig. 1). The phylogenetic distribution of our newly sequenced genomes indicates a high diversity of soil-dwelling *Bradyrhizobium*.

Our ancestral lifestyle reconstruction analysis inferred that the last common ancestor (LCA) of *Bradyrhizobium* adapted to a free-living lifestyle, consistent with our prior study based on a very limited taxon sampling [9] (Fig. 1). This was evident by the pattern that a vast majority of the outgroup lineages as well as all members of the basal *Bradyrhizobium* clade (Soil 1 and Soil 2 supergroups) were non-symbiotic lineages. The symbiotic lifestyle independently originated eight times based on the sampled lineages, including three major origins: one in the *B. jicamae* supergroup (FL_nonnif_-Sym1), one in the *B. elkanii* supergroup (FL_nonnif_-Sym3), and another one at the LCA of the Kakadu, Photosynthetic, and *B. japonicum* supergroups (FL_nonnif_-Sym4). These transitions occurred in relatively deep phylogenetic positions, suggesting that they represent evolutionarily ancient events. Free-living lifestyles originated from symbiotic strains 23 times: 13 of these events gave rise to non-N_2_-fixing strains and 10 gave rise to N_2_-fixing members (Fig. 1). Transitions from symbiotic to free-living lifestyle (both Sym-FL_nif_ and Sym-FL_nonnif_) likely occurred recently as indicated by their shallow phylogenetic positions (Fig. 1). Hence, the loss of the ability to nodulate legume plants occurred more frequently than the reverse process in *Bradyrhizobium*, consistent with a prior study based on the analysis of phenotypic markers [7]. This pattern remained when we combined FL_nif_ and FL_nonnif_ as a single free-living lifestyle (Fig. S2). Furthermore, the free-living N_2_-fixing lifestyle originated from free-living non-N_2_-fixing ancestors four times (Fig. 1). The infrequency of this type of lifestyle transition does not necessarily mean that this type of transition is rare, as it may result from a potential bacterial cultivation bias against those carrying the *nif* cluster under aerobic conditions with readily available N sources.

### HGT of the *nif* island drives the expansion of the free-living N_2_-fixing lifestyle

Free-living N_2_-fixing *Bradyrhizobium* is of much interest, because these *Bradyrhizobium* members may play a previously unrecognized role in the N cycle in soils, especially given the reportedly high abundance of *Bradyrhizobium* in soils [6]. We asked where the N_2_-fixing (*nif*) genes in free-living *Bradyrhizobium* come from and whether the *nif* genes differ between free-living and symbiotic members. Since most free-living strains evolved from their symbiotic ancestors, one possibility is that free-living N_2_-fixing strains inherited the same version of *nif* genes from their nodulating ancestors. Alternatively, free-living N_2_-fixing *Bradyrhizobium* might have lost the entire symbiosis island which includes both *nod* and *nif* clusters among other genes, and instead recruited a new set of *nif* genes from an external donor by HGT. To test these competing hypotheses, we constructed the phylogenetic tree of the *nif* cluster based on the concatenated alignment of the genes *nifABDEHKNVX* for those from the *Bradyrhizobium* and several other rhizobia including *Azorhizobium*, *Rhizobium*, *Sinorhizobium* and *Mesorhizobium* from *Alphaproteobacteria*, and *Burkholderia* and *Cupriavidus* from *Betaproteobacteria* (see Materials and methods).

The *nif* genes of all *Bradyrhizobium* formed a monophyletic group (Fig. S3), indicating a single origin of *nif* in this genus. Within the *Bradyrhizobium* clade, the *nif* gene tree (Fig. 2) displayed substantial topological incongruence with the species tree (Fig. 1), suggesting different evolutionary histories between *nif* genes and the core genome. In the *Bradyrhizobium* part of the *nif* gene tree, the Photosynthetic *Bradyrhizobium* supergroup together with a strain from the *B. jicamae* supergroup (SZCCT0283) branched first, followed by a few strains of the *B. japonicum* supergroup (Fig. 2). The latter consisted of two clades, one exclusively constituted by symbiotic strains (SymBasal; Fig. 2), some of which were early-split lineages of the *B. japonicum* supergroup (Fig. 1), and the other mostly composed of free-living N_2_-fixing members (Cluster FL) (Fig. 2). *Nif* genes of the *B. jicamae* and *B. elkanii* supergroups, which together with *B. japonicum* comprised the three largest clades in the *nif* gene tree (Fig. 2), were nested within the *B. japonicum* supergroup, suggesting that they were horizontally transferred from the latter. Despite much inconsistency between the species and *nif* trees, but because the basal positions in the *nif* phylogeny mostly included members from Photosynthetic, Kakadu, and *B. japonicum* supergroups, it is likely that the *nif* cluster in *Bradyrhizobium* was first acquired by the LCA of the Photosynthetic, Kakadu and *B. japonicum* supergroups. This is consistent with the ancestral lifestyle reconstruction shown in Fig. 1 which inferred a symbiotic ancestor of the corresponding ancestral node (FL_nonnif_-Sym4).

Strikingly, despite a scattered distribution in the species phylogeny (Fig. 1), 17 free-living N_2_-fixing strains grouped on the *nif* phylogeny (Cluster FL in Fig. 2). Also grouping in Cluster FL were the *nif* genes from three symbiotic strains (DOA9, CCBAU 53363 and p9-20). These three strains encoded another *nif* cluster located in the symbiotic plasmid (DOA9) or symbiosis island on the chromosome (CCBAU 53363 and p9-20), which grouped with other symbiotic strains in the *nif* phylogeny (Fig. 2). Clearly, the grouping of the *nif* genes belonging to Cluster FL was the result of HGT. This point was further evidenced by the conserved genomic context around *nif* genes in Cluster FL (see the next section). Moreover, the geographic locations of isolates belonging to Cluster FL spans three continents (North America, South America and Asia) (Fig. 1B). Remarkably, as introduced above, two strains (*Bradyrhizobium* sp. AT1 and *Bradyrhizobium* sp. DOA9) whose *nif* genes from Cluster FL in the *nif* phylogeny were previously shown to perform N_2_ fixation under micro-aerobic conditions (marked by a box around the strain name in Fig. 2) [20, 21]. The above evidence together indicates that *nif* genes from Cluster FL, which were mainly comprised by free-living *Bradyrhizobium*, are likely to be a special group of *nif* genes that specifically take part in free-living N_2_ fixation.

### A unique *nif* island likely contributes to the oxygen tolerance of free-living N_2_-fixing *Bradyrhizobium*

We sought more evidence for HGT of the *nif* cluster by analyzing genes surrounding it. This led to the identification of a ~50 kb genomic region containing *nif* genes (*nif* island) conserved among all members of Cluster FL (Fig. 3A). The regions flanking the *nif* island were not conserved across most free-living strains (Fig. S4), providing further evidence for HGT of the *nif* island. Note that four strains of Cluster FL (BM-T, Y-H1, R2.2-H and W) shared highly conserved genomic contexts (Fig. S4). These strains were closely related and comprised a monophyletic group in the phylogenomic tree (marked by a box around the strain names in Fig. S5), indicating that the shared genomic context of their *nif* islands was inherited from their LCA. Additionally, the conserved genomic context of the *nif* island among strains from the Photosynthetic *Bradyrhizobium* supergroup (Fig. S4) was consistent with their vertical descent shown in the species phylogeny (Fig. 1). We further compared this unique *nif* island with the *nif* clusters of other strains in this genus (Fig. 3).

**Figure 3.**
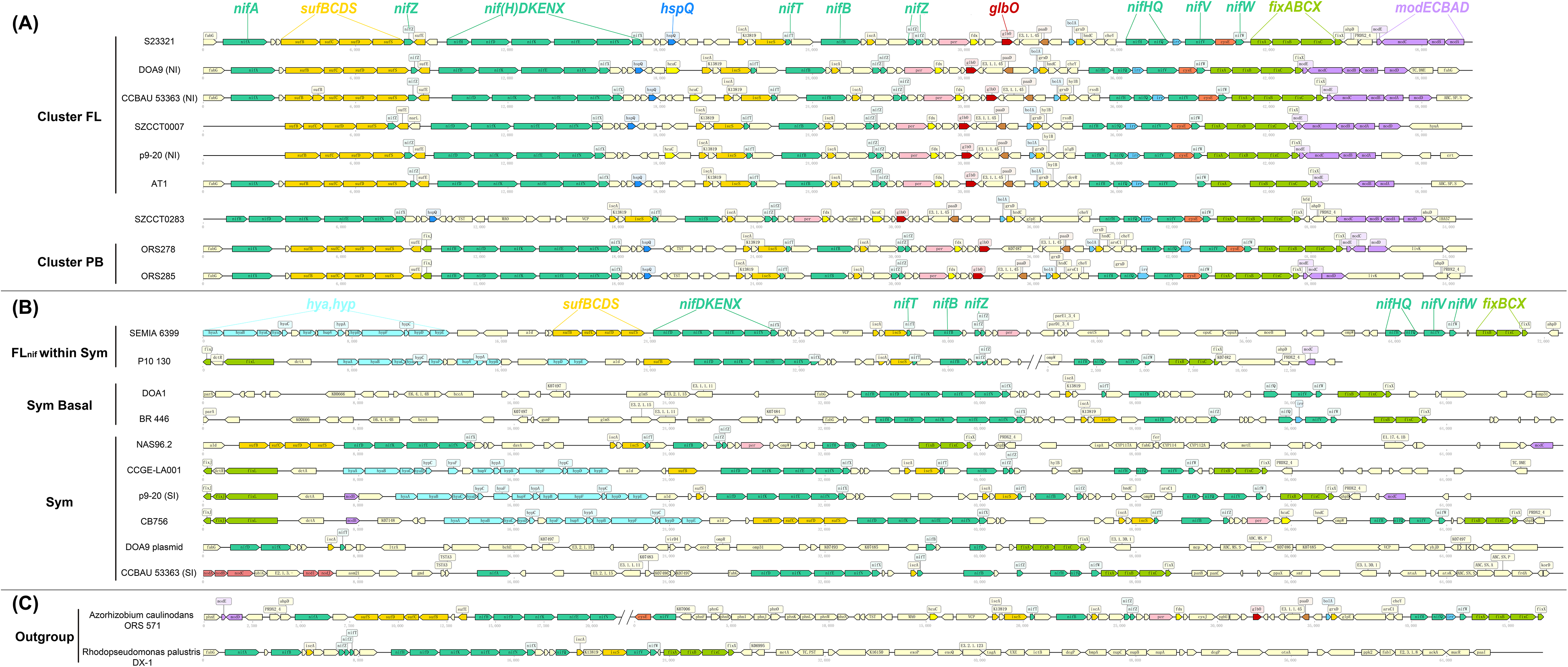
The comparison of the gene arrangement of the *nif* gene cluster located in the *nif* island and in the symbiosis island. Gene functions are distinguished by different colors. The visualization of gene arrangement is performed with dna-features-viewer v3.0.3 [64]. The structures of the gene arrangement classified the *nif* clusters into three types: those belonging to the Cluster FL or Cluster PB (A), those found in the symbiosis island in most symbiotic strains or in FL_nif_ strains that cluster with symbiotic strains (B), and those from outgroup species (C). In panel B, two out of the three free-living strains whose phylogenetic positions are nested within symbiotic strains, labeled as “FL_nif_withinSym”, are shown together with the symbiotic strains closest to them in the *nif* tree (see also Fig. 2): SEMIA 6399 vs. NAS96.2, and CCGE-LA001 vs. P10 130 (the other one, strain CCBAU 83689, is not shown as the assembly of the contig where the *nif* island is located is incomplete).

The upstream boundary of the *nif* island in Cluster FL *Bradyrhizobium* (Fig. 3A) was marked by *nifA*, which codes for a transcriptional activator required for the expression of *nif* operons [34]. Adjacent to *nifA* was a *suf* cluster, which is responsible for synthesizing and inserting Fe-S clusters into nitrogenase [35]. The core genes encoding the nitrogenase were located downstream of the *suf* cluster, and were mainly separated into two operons, *nifDKENX* and *nifHQ*. *FixABCX* which encode a membrane complex involved in electron transfer to nitrogenase were located downstream of the *nif* genes [36]. Extensively studied in rhizobia, the role of *fixABCX* in N_2_ fixation has been suggested for free-living diazotrophs [37]. At the downstream boundary of the *nif* island was a *mod* cluster encoding molybdate transporter.

In general, the *nif* island conserved in free-living *Bradyrhizobium* was similar to that in photosynthetic strains (Fig. 3A), as also noted in previous studies [38, 39], and that in a newly sequenced soil-dwelling strain (SZCCT0283) from the *B. jicamae* supergroup (Fig. 3A), consistent with the evolutionary relatedness of these *nif* islands (Fig. 2). A few differences were that those from Cluster PB carried an additional copy of *nifH* but fewer genes in the *mod* operon compared with those from Cluster FL (Fig. 3). Further, the early-split positions of Cluster FL and PB *nif* genes in the phylogeny (Fig. 2) suggest an ancient evolutionary origin. Because *nif* genes in the symbiosis island of most symbiotic strains branched later than those from the *nif* island (Fig. 2), presumably, the *nif* island found in photosynthetic and free-living *Bradyrhizobium* represent the ancestral status of symbiotic *Bradyrhizobium*, and it was later expanded to a symbiosis island by integrating *nod* as well as other genes in non-photosynthetic symbiotic *Bradyrhizobium*.

The *nif* clusters of the *nif* island conserved in Cluster PB and FL (Fig. 3A) shared many genes with those on the symbiosis island (Fig. 3B), such as *nifDKENBH* and *fixBCX*, agreeing with their essential roles in N_2_ fixation. However, they also displayed notable differences. Apparently, the gene arrangement of the free-living *nif* island was more compact and more conserved across different strains (Fig. 3). In contrast, in the *nif* cluster from the symbiosis island, several genes involved in N_2_ fixation like *nifA* and *fixA* are often linked to *nod* genes and separated from the *nif* cluster [40]. The *nif* island in free-living and photosynthetic members additionally carried a *nifV* gene, which encodes homocitrate synthase to synthesize homocitrate, a component of the Fe-Mo cofactor of nitrogenase [41]. The vast majority of symbiotic rhizobia (not limited to *Bradyrhizobium*) do not harbor *nifV* [41], which is known to be compensated for by the homocitrate provided by their legume host during the symbiotic state [42]. Some studies attributed the ability of free-living N_2_ fixation to *nifV*, because this gene was detected in the genomes of the few rhizobial strains able to perform N_2_ fixation in the free-living state [41, 43, 44]. Here, we found that *nifV* was present in around half of *Bradyrhizobium* genomes, not restricted to the free-living and photosynthetic strains (Fig. 2). Phylogenetic analysis suggests that *nifV* was presumably present at the LCA of *Bradyrhizobium*, but lost within the *B. japonicum* supergroup (Node 1 in Fig. 2; Fig. S6). Further, early studies showed that *Bradyrhizobium* sp. CB756, a strain carrying *nifV* on the symbiosis island (Fig. 3B), fixed N_2_ at a rate more than a magnitude lower than the *nif* island-carrying photosynthetic *Bradyrhizobium* (strains ORS310 and ORS322) and *Azorhizobium caulinodans* ORS571 in agar culture under air or O_2_ concentrations resembling soil environments [19, 45]. This hints that the absence of *nifV* might not be the only reason accounting for the decreased efficiency of free-living N_2_ fixation of certain rhizobia strains, at least at higher O_2_ concentrations.

We further identified that several genes involved in O_2_ tolerance and stress response were also specific to the *nif* island of free-living and photosynthetic members (Fig. 3), including *glbO*, *hspQ*, and the *mod* operon. Specifically, *glbO* encodes a 17-kDa group-II truncated hemoglobin that binds O_2_ with high affinity [46, 47]. The expression of truncated hemoglobin genes is induced upon exposure to oxidative stress generated by H_2_O_2_ and hypoxia [48]. The product of *glbO* might therefore play an important role in protecting nitrogenase from O_2_ inactivation, endowing the free-living *Bradyrhizobium* with higher tolerance to O_2_. *HspQ* encodes a chaperone protein, which combats the detrimental effects on proteins caused by stressors such as increased temperatures, oxidative stress, and heavy metals [49]. The *mod* operon transports extracellular molybdate with high affinity [50]. We also observed copy number differences between the free-living and symbiotic members, such as the *nif*Z which has a function in the maturation of the Fe-Mo protein nitrogenase [51], but whether the additional copies contribute to adaptation to the free-living lifestyle is still unknown. In summary, we predict that in addition to *nifV*, the genes involved in oxygen tolerance and stress response in the *nif* island (e.g., *glbO*, *hspQ* and *mod*) may play a role in the adaptation to free-living lifestyle. Intriguingly, the *nif* island from *Azorhizobium*, which is able to perform free-living N_2_ fixation, shared some of the above genes like *nifV*, *glbO* and *mod*, and a generally similar gene arrangement with the free-living *nif* island in *Bradyrhizobium* (Fig. 3C).

Note that, for simplicity, we defined free-living strains based on the absence of *nod* in the current study. Four free-living N_2_-fixing strains, namely *B. mercantei* SEMIA 6399, *B. yuanmingense* P10 130, *B. liaoningense* CCBAU 83689 and *B. mercantei* SEMIA 6399, were originally isolated from nodules (Data Set S2). Interestingly, except the last isolate whose *nif* genes were found in Cluster FL, the *nif* genes of the other three were embedded in symbiotic lineages in the *nif* phylogeny (Fig. 2), indicating that the *nif* genes were inherited from nodulating ancestors. These three strains were also found to carry genes encoding components of T3SS and the effectors it injects, as evident by the genes gained or lost shown in Fig. S7C (Sym-FL_nonnif_ transitions 2, 5 and 6; Text S2), suggesting that they are likely nodule-inhabiting strains that do not nodulate but with the potential to interact with plants. This is in contrast to other free-living N_2_-fixing strains where symbiosis genes were commonly lost during lifestyle transition (Sym-FL_nonnif_ transitions 1, 3, 4 and 7-10 in Fig. S7C; Text S2). In general, the gene arrangement of *nif* genes and surrounding regions for these strains were similar to those of symbiotic strains, particularly when compared with their closest relatives, but the degree of similarity varied: while P10 130 showed a very high similarity to its closest *nod*-carrying relative CCGE-LA001 in the genomic context of the *nif* genes, SEMIA 6399 showed a limited similarity to its closest *nod*-carrying relative NAS96.2, likely due to a recent insertion in SEMIA 6399 of several genes between *nifZ* and *nifH* (Fig. 3B). We did not analyze CCBAU 83689 as the contig carrying *nif* genes was too short. The results indicate the different origins of the free-living N_2_-fixing *Bradyrhizobium*: while some were derived from their symbiotic ancestors, most analyzed strains of this lifestyle originated via HGT of the free-living *nif* island (Fig. 3). It would be interesting to conduct experiments to assess the detailed functions of the genes on the *nif* island under O_2_ concentrations resembling those of soil environments in future.

### Widespread distribution of the unique free-living nif island in the environment

We further asked how prevalent the *nif* island specific to free-living *Bradyrhizobium* is in the environments and how it compares to the *nif* genes of symbiotic members. We therefore assessed the normalized abundances of *nifH*, the most commonly used marker gene to identify N_2_-fixing microbes [52], that potentially correspond to free-living (Cluster FL in Fig. 2), photosynthetic (Cluster PB in Fig. 2), and symbiotic *Bradyrhizobium* (Sym and SymBasal in Fig. 2) in 4,958 amplicon sequencing data sets (see Materials and methods). We divided these data sets into four categories according to the environments where the samples were collected: soils (e.g., bulk soil, rhizosphere and freshwater lake sediment), plants (root and phyllosphere samples), marine (including seawater, marine sediment, coral and mangrove samples), and others (e.g., bioreactor and unidentified samples) (Data Set S3). Amplicon reads were assigned to different types of *Bradyrhizobium nifH* according to their phylogenetic placements (see Materials and methods).

Amplicon analysis showed that *Bradyrhizobium* constitute an important group of diazotrophic communities in different habitats, and the relative abundance of *Bradyrhizobium* was 7.52±18.57%, 6.67±12.75%, 17.39±18.32, 14.54±18.85% in marine, soil, plant, and other environmental types, respectively (Data Set S3). The *nifH* that resembles those belonging to Cluster FL displayed the highest normalized abundance in samples from marine, soil, and habitats classified as “others” (Fig. 4; *p* < 0.001 for all comparisons, paired Wilcoxon-Mann-Whitney test). Specifically, Cluster FL accounted for 56±47%, 46±38%, and 51 ±39% (mean±standard deviation) of reads that were assigned to *Bradyrhizobium* in these three types of habitats (marine, soil, and others), respectively. In particular, for marine samples, nearly all *Bradyrhizobium nifH* were assigned to Cluster FL (Data Set S3). For the data sets of plant samples, while symbiotic *nifH* displayed the highest abundance (Fig. 4), *nifH* assigned to Cluster FL also showed a considerably high normalized abundance in plants (29±39%), and was even the dominant ecotype in diazotrophic communities associated with some Asteraceae species and the roots of several perennial grass species (Data Set S3). Hence, though the presence of free-living *Bradyrhizobium*-specific *nifH* does not necessarily indicate the occurrence of a complete *nif* island, it is tempting to suggest that *Bradyrhizobium* members carrying the free-living *nif* island i) were distributed in a variety of environments (Fig. S8), and ii) were in general more abundant than that of symbiotic strains in the genus in most habitats except plants, hinting at a previously unrecognized role of their *nif* island in the free-living lifestyle. Nevertheless, it should be noted that Cluster FL *nifH* also displayed large variations in regard to the normalized abundance between different diazotrophic communities of the same type of habitat despite the general trend described above.

**Figure 4.**
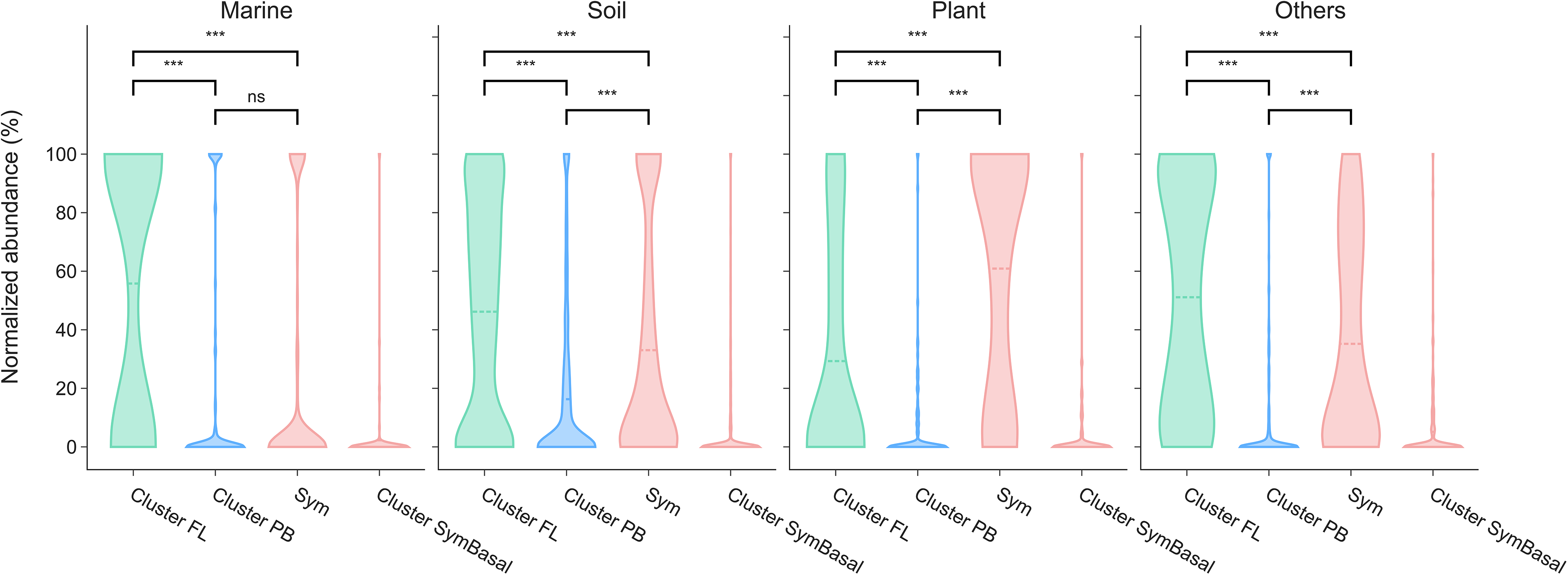
Normalized abundance of free-living and symbiotic *nifH* of *Bradyrhizobium* in amplicon sequencing samples collected from different types of environments. The values shown as percentage in the y-axis in the violin plots denote the normalized abundance (see Materials and methods) of different types of *nifH*. The width of each curve represents the approximate frequency of data points in the corresponding region. The *P*-value is derived from a paired Wilcoxon-Mann-Whitney test (two-sided).

## Caveats and concluding remarks

Adding the 93 soil-dwelling strains greatly expanded our knowledge of the ecology and evolution of *Bradyrhizobium*. However, because of the bias towards symbiotic strains in previous genomics research and the difficulties in cultivating those dwelling soil environments, the free-living strains analyzed here may underestimate the real diversity of wild diazotrophic *Bradyrhizobium* that adapt to a free-living lifestyle. For example, prior studies have shown that *Bradyrhizobium* is the dominant genus in the diazotrophic communities in many soil ecosystems, most of which have not been sampled for *Bradyrhizobium* cultivation however [53, 54]. Of the limited types of soil ecosystems sampled here, the *Bradyrhizobium* strains we collected are not necessarily representative of the wild populations in these ecosystems owing to the potential cultivation bias. For example, our cultivation process was under aerobic condition and the cultivation medium was rich in fixed nitrogen, conditions disfavoring the isolation of diazotrophic members. Therefore, it is not clear whether important free-living diazotrophic lineages occupying distinct phylogenetic positions are missing and how this complements the current conclusion we reached. Another caveat to some of our findings is that some of our analyses built on an over-simplified classification of *Bradyrhizobium* lifestyles, which was based on the presence/absence of certain genes like *nif* and *nod*. In fact, isolates with a symbiosis island could be ineffective (i.e., those form nodules but fix little N_2_ within nodules) [55]. Likewise, whether the free-living strains possessing the *nif* island could perform N_2_ fixation, and if they could, the N_2_ fixation efficiency under different O_2_ concentrations, remains to be studied.

In spite of the above caveats, our study reveals an interesting pattern of convergent evolution of free-living N_2_-fixing *Bradyrhizobium* from their symbiotic ancestors. Though the nested phylogenetic positions within symbiotic lineages (Fig. 1) might leave an impression that free-living N_2_-fixing members could have inherited the *nif* genes from their symbiotic ancestors, we provided compelling evidence that this is unlikely the case. Rather, it is the HGT of a conserved *nif* island, which presumably represents the ancestral state of all types of *nif* found in the *Bradyrhizobium*, that drives the independent transitions from symbiotic to free-living N_2_-fixing lineages. This serves as a classic example of convergent shifts in lifestyle driven by HGT of certain genes in bacteria, which, although mostly explored in pathogens or symbionts in prior studies [56–58], may play a prominent role in free-living bacteria. It further hints that the symbiosis island common to the vast majority of symbiotic *Bradyrhizobium* members could be derived from the ancestral *nif* island potentially by recruiting symbiosis genes. This may be different from the traditional view that the symbiosis island, including *nif*, *nod* and T3SS among other genes, was acquired in its entirety to enable ancestral interactions between *Bradyrhizobium* and legumes [9, 59]. Given the global dominance of *Bradyrhizobium* in the soil microbiota, even a small proportion of them being diazotrophic members could potentially bring a considerable amount of fixed nitrogen to the bulk soils and other nitrogen-limited terrestrial ecosystems. Our results therefore have important implications for understanding terrestrial nitrogen cycles. Because one of the major aims of modern agricultural research is to transfer the N_2_ fixation ability to non-legume crops [60, 61], the capacity of certain free-living *Bradyrhizobium* to fix N_2_ and their association with non-leguminous plants may imply potential application in agriculture.

## Acknowledgements

We thank Xiaoyuan Feng for help with genome assembly. We appreciate Shuang Wang and Jie Liu for providing soil samples and Jianjun Xu for assisting in sample collection. We are grateful to Xingqin Lin for her helpful suggestions in medium design. We also thank Qin Li, Yan Li, Xiaojun Wang, Xiao Chu, Hao Zhang and Danli Luo for their helpful discussion. This work was supported by the Hong Kong Research Grants Council Area of Excellence Scheme(AoE/M-403/16).

## Data and codes availability

The genomic sequences of the 88 sequenced *Bradyrhizobium* and five *Afipia* strains have been submitted to the NCBI databases GenBank (accessions: PRJNA698083), which will be made publicly available upon the acceptance of the manuscript. The Python [62] and Ruby [63] codes used for phylogenomic and comparative genomics analyses are deposited in the online repository https://github.com/luolab-cuhk/Bradyrhizobium-Nif-HGT.

