## Supplementary information for "Horizontal gene transfer of a unique *nif* island drives convergent evolution of free-living N_2_-fixing *Bradyrhizobium*"

**This file includes:**

Supplementary Text S1. Supplementary methods

Supplementary Text S2. Other metabolic strategies facilitating repeated ecological expansions from legume hosts to soil environments

References

Figures S1 to S8

### Supplementary Text 1

#### 1.1 Supplementary methods

##### Soil and plant tissue sampling and processing

Samples were collected from five different sites in China (Fig. S1A) that cover several ecosystem types: soybean cropland (Heihe and Lvliang), artificial park (Shenzhen), undeveloped forest (Lanzhou), and grassland (Hefei). In Heihe, Lvliang and Hefei sites, intact *Glycine max* or *Erigeron annuus* plants were excavated from soil, and shaken mildly to remove soil loosely adhering to the roots. The remaining adherent soil was separated from the roots as rhizosphere soil. For Shenzhen and Lanzhou sites, 0-5 cm and 5-10 cm depth bulk soil was collected from the vicinity of plant root. At the Shenzhen sampling site, the soil was collected near *Acacia confusa* (legumes), *Calliandra haematocephala* (legumes) and *Bambusoideae* (non-legumes). At the Lanzhou site, the soil was collected near non-leguminous plant *Picea asperata*. Detailed information about sampling sites was provided in Data Set S1. Soil samples were placed in sterile bags, kept on ice, and immediately transported to the laboratory for further processing.

##### *Bradyrhizobium* isolation and identification

To prepare soil inoculum, 5.0 g of fresh soil was put in a 50 mL conical tube with 45 mL of sterile deionized water. After mixing with a vortex mixer, 1 mL of soil suspension was serially diluted, and the dilutions were used for inoculation. Roots of *Erigeron annuus* were processed according to Coombs and Franco [1]. One gram of surface-sterilized roots was grinded in a mortar and diluted with PBS buffer, and the dilutions of the root slurry were used for inoculation.

Five different media were applied to retrieve target strains from the samples, including modified arabinose-gluconate (MAG) media adapted from the study Sachs *et al* [2], 10- and 100-fold diluted MAG, ρMAG (MAG supplemented with ρ-coumaric acid) and vanillic acid media (detailed in Supplementary Text 1.1). All media were supplemented with 55 mg/L cycloheximide to inhibit fungal growth. All agar plates were incubated at 28 °C . Colonies with small, white and raised morphology formed after 7 days were picked for species identification.

Colonies were identified by a 1,465 bp fragment PCR product using universal bacteria 16S rRNA primers 27F and 1492R. Chelex 100 resin [Bio-rad, USA] was used to extract DNA from bacterial colonies for PCR reaction, and the recipe of PCR was prepared using Premix Taq [Takara Bio, USA]. The PCR conditions were as follows: denaturation at 95℃ for 5 minutes, followed by 32 cycles (95℃ for 45 seconds, 55℃ for 45 seconds and 72℃ for 90 seconds), final extension at 72℃ for 10 minutes. The taxonomic information of the isolates was obtained by comparing the 16S rRNA gene sequences using EzBioCloud [3], which is a taxonomically united database of 16S rRNA gene sequences and whole-genome assemblies.

**Genomic DNA extraction and sequencing**

Genomic DNA was extracted using OMEGA Bacteria DNA kit (D3350-02, OMEGA Bio-tek, USA) according to the manufacturer's instructions. The quantity of extracted DNA was evaluated using a Nanodrop ND-1000 spectrophotometer (Nanodrop Technologies, Wilmington, DE) and 0.8% agarose electrophoresis. Whole genome sequencing of the isolates was performed using paired-end sequencing method with Hiseq X platform (Illumina) at Magigene Company (Guangzhou, China). After DNA extraction and sequencing, raw reads were quality trimmed with Trimmomatic v0.36 [4] with the options 'SLIDINGWINDOW:4:15 MAXINFO:40:0.9

MINLEN:40' and assembled using SPAdes v3.10.1 [5] with the option '-careful'. Only contigs with length > 1,000 bp and sequencing depth ≥ 5 were retained. Protein-encoding genes were predicted with Prokka v1.12 [6]. Completeness of these genomes was calculated using CheckM v1.0.7 with default parameters.

Protein-coding genes were annotated by searching against the KEGG hmm profiles (2019 release) [7] using HMMER 3.2.1 [8] with an e-value cutoff of 1e-20. We further searched genes not annotated by KEGG against COG [9] and TIGRFAM [10] databases using diamond v0.9.3.104 [11] with the e-value cutoff of 1e-20.

**Phylogenetic analysis**

We downloaded 258 reference genomes from the NCBI GenBank (July, 2019), including 247 genomes belonging to *Bradyrhizobiaceae* family and 11 from other *Rhizobiales* lineages as the outgroup (Data Set S2). Six strains that were either wrongly assigned to *Bradyrhizobium* (CCH5-A9, CCH1-B1 and HPC_L) or had a genome completeness lower than 50% (JGI_0001019-M21, JGI_0001019-J21 and JGI_0001002-A22) were excluded. The remaining 252 genomes, together with the 93 *Bradyrhizobium* strains sequenced in the present study, were used for phylogenomic construction based on 123 shared single-copy genes identified by OrthoFinder v2.2.7 [12]. Genes were aligned at the amino acid level using MAFFT v7.222 [13] and the alignments were trimmed with trimAl v1.4 which removed sites with a similarity score lower than 0.001 ("-st 0.001") [14]. The maximum likelihood (ML) phylogenomic tree was constructed based on the concatenated alignment of these proteins using IQ-Tree v1.6.2 [15] with the parameters "-s alignment -spp partition -m MFP+MERGE -rcluster 10 -bb 1000 -wbtl -mset WAG,LG,JTT -mrate E,G,I,G+I", which finds the best partitioning scheme using the edgeproportional partition model for the concatenated alignment and allows each partition to have its own substitution models [16]. Node supports were assessed by performing 1,000 ultrafast bootstrap replicates, and only those with support value ≥ 95% were considered to be reliable [17]. All trees were edited and displayed with iTOL v4 (https://itol.embl.de/).

**Reconstructing gene family evolution**

The ancestral gene gains and losses of all protein families along the phylogenomic tree was reconstructed using BadiRate v1.35 [18] with the parameter " --ep CWP --anc -rmodel BDI -bmodel FR". Gene family turnover rates were estimated using CWP (Wagner parsimony) algorithm under the BDI (Birth, Death and Innovation) and FR branch model (Free Rates), which assumes that each branch in the phylogenomic tree has its own turnover rate.

**Comparing the relative abundances of diazotrophic *Bradyrhizobium* displaying different lifestyles in the environments**

Metadata of 4,958 sequence files (SRA runs) were retrieved from the NCBI Sequence Read Archive (SRA) using the term "*nifH* metagenome" (last accessed in December 2020). We downloaded raw data using SRA Toolkit (https://github.com/ncbi/sra-tools). Quality control was performed by Trimmomatic v0.39 [4] with default parameters (LEADING and TRAILING:3; MINLEN:50 SLIDINGWINDOW:4:15). For sequence files with base quality information (Phred quality score), we employed DADA2 [19] implemented in QIIME2 [20] to denoise sequences (removal of noisy sequences, chimeric sequences and singletons, joining denoised paired-end reads, and dereplication). For those without quality information, VSEARCH [21] pipelines from

QIIME2 were used to cluster sequences at an identity cutoff of 97%.

We applied evolutionary placement methods to assign the short sequence reads to the four phylogenetically distinguishable types of *nif* clusters (i.e., FL, PB, Sym and SymBasal; see Fig. 2) based on the following procedure. We first aligned reference *nifH* genes used in Fig. 2 and in Wang *et al* [22] using the GINSI algorithm implemented in MAFFT v7.222 [13]. This alignment served as a backbone to which we aligned short sequence reads from amplicon sequencing data sets using PaPaRa v2.5 with the default parameters [23]. Next, we generated a phylogeny of the reference *nifH* genes using IQ-Tree v1.6.2 [15] with the parameters "-s alignment -m MFP -wbtl -bb 1000 -mset GTR,HKY85,K80". Finally, we performed phylogenetic placements using EPA-ng v0.3.8 [24] with the parameters "GTR{0.7/1.8/1.2/0.6/3.0/1.0}+F+R10" whose values were estimated by IQ-Tree. The normalized abundance of *nifH* genes classified to each type of *nifH* genes was calculated as the number of reads assigned to the corresponding *nifH* type divided by the total number of reads assigned to *Bradyrhizobium* in the amplicon sequencing data set. Note that SZCCT0283 was excluded in the analysis because it could belong to none of the four types of *nifH* based on its position in the *nif* phylogeny (Fig. 2).

**1.2 Medium recipe**

**MAG (Modified Arabinose-Gluconate) Media Reagents**

1.1g MES [2-(*N*-morpholino)ethanesulfonic acid]

1.3g HEPES [4-(2-hydroxyethyl)-1-piperazineethanesulfonic acid]

1.0g Arabinose (DL)

1.0g Sodium gluconate

1.0g Yeast extract

2ml $KH_2PO_4$ Solution (110g/L)

4ml $Na_2SO_4$ Solution (62.5 g/L)

1ml $MgSO_4 \cdot 7H_2O$ Solution (180g/L)

2ml $NH_4Cl$ Solution (160g/L)

1ml $CaCl_2$ Solution (13g/L)

1ml $FeCl_3 \cdot 6H_2O$ Solution (6.7g/L)

Adjust pH to 6.6 with KOH

Add 15 g Agar and replenish the liquid to 1L

**ρMAG Media Reagents**

1.1g MES

1.3g HEPES

0.1g ρ-cumaric acid

1.0g Arabinose (DL)

1.0g Sodium gluconate

1.0g Yeast extract

2ml $KH_2PO_4$ Solution (110g/L)

4ml $Na_2SO_4$ Solution (62.5 g/L)

1ml $MgSO_4 \cdot 7H_2O$ Solution (180g/L)

2ml $NH_4Cl$ Solution (160g/L)

1ml $CaCl_2$ Solution (13g/L)

1ml $FeCl_3 \cdot 6H_2O$ Solution (6.7g/L)

Adjust pH to 6.6 with KOH

Add 15 g Agar and replenish the liquid to 1L

**V 0.1 Media Reagents**

0.11g MES

0.13 HEPES

0.1g Vanillic acid

100µl $KH_2PO_4$ Solution (110g/L)

200µl $Na_2SO_4$ Solution (62.5 g/L)

500µl $MgSO_4 \cdot 7H_2O$ Solution (180g/L)

100µl $NH_4Cl$ Solution (160g/L)

50µl $CaCl_2$ Solution (13g/L)

50µl $FeCl_3 \cdot 6H_2O$ Solution (6.7g/L)

Adjust pH to 6.6 with KOH

Add 7.5 g Agar and replenish the liquid to 500 ml

**Supplementary Text S2. Other metabolic strategies facilitating repeated ecological expansions from legume hosts to soil environments**

To further understand the genomic changes during the transition from symbiotic to free-living lifestyle, we reconstructed the genome content at ancestral nodes using BadiRate, and inferred the gains/losses of genes during the process of this type of lifestyle transition (see Materials and methods) (Fig. S7; Data Sets S4-S6). Previous genomics studies in rhizobia [22, 25, 26] mostly focus on the evolution towards the symbiotic lifestyle (Fig. S7A), leaving the genomic changes associated with the transitions from symbiotic to free-living lifestyle largely unknown. In the present study, we focused on Sym-$FL_{nonnif}$. We found that this type of lifestyle transition was characterized by losses of myriad symbiosis-related genes, most of which are located in the symbiosis island (Fig. S7B). Genes repeatedly lost during this type of transition included those for nodulation (*nod*, *nol* and *noe*), $N_2$ fixation (*nif*) and its regulation (*fix* and *hya*), and Type III (T3SS) and IV (T4SS) secretion systems which inject effectors into host cells to establish symbiosis (Fig. S7B). Previous studies mainly focused on T3SS as a main secretion system in the establishment of nodulation and determining host specificity [27]. Our results agree with this but additionally hint at the role of the (*trb* type) T4SS in legume-*Bradyrhizobium* symbiosis. Many genes encoding transposable elements were lost when a symbiotic strain was converted to a free-living strain, possibly a result of the loss of symbiosis island where many transposons are located [28]. Also repeatedly lost during Sym-$FL_{nonnif}$ were genes involved in cellular interaction (e.g., the quorum sensing system regulator *CciR*), biotin synthesis, and bacterial growth in nodules [29, 30] (Fig. S7B).

Different from Sym-$FL_{nonnif}$, Sym-$FL_{nif}$ did not show copy number changes in $N_2$-fixing related genes *nif* and *fix* (Fig. S7C): as mentioned above, they acquired a free-living *nif* island after losing the symbiosis island. Also with no changes in copy numbers during this type of lifestyle transition were hydrogenase genes (*hya*; Fig. S7C). Although genes encoding hydrogenase are often located in the symbiosis island in symbiotic strains (Fig. 3) [31, 32], they were absent in the *nif* island in free-living strains (Fig. 3A). Thus, the presence of these genes in free-living $N_2$ fixing strains should not be the result of vertical inheritance as the entire symbiosis island in their symbiotic ancestors is supposed to be removed during lifestyle transition. Presumably, they were gained during lifestyle transition by HGT independent of the *nif* island (Fig. S7C). This pattern suggests that hydrogenase may be beneficial to $N_2$ fixation (in the free-living state), consistent with the previous experimental results [33].

Most genes that were likely acquired during the transitions from symbiotic to free-living lifestyle were found to occur in a lineage-specific manner. This may be illustrated by genes involved in denitrification (Data Set S6): while the cytoplasmic $NO_3^-$ reductase *nar*, *nir*, *nor* and *nos* genes were acquired in some lineages, in other lineages their copy numbers remained unchanged (Fig. S7C). In general, we did not identify genes that displayed a clear pattern of repeated gains during the transition from symbiotic ancestors. This result, however, is not surprising. This is because symbiotic rhizobia do not always live in nodules. When they are released from nodules they have to survive the soil environments. Clearly, they are supposed to be equipped with genes helping them dwell nodules, rhizosphere, and soils [22]. Due in part to this reason, *Bradyrhizobium* are known to have some of the largest genomes in bacteria [34]. This is also consistent with the observations that the transition from symbiotic to free-living lifestyle was easier than the opposite process since losses of genes involved in symbiosis may be more likely to happen than gains of them [22].

28. Kaneko T, Nakamura Y, Sato S, Minamisawa K, Uchiumi T, Sasamoto S, et al. Complete genomic sequence of nitrogen-fixing symbiotic bacterium Bradyrhizobium japonicum

USDA110. *DNA Research* 2002; **9:** 189-197.

34. Ormeno-Orrillo E, Martinez-Romero E. A genomotaxonomy view of the Bradyrhizobium genus. *Frontiers in microbiology* 2019; **10:** 1334.

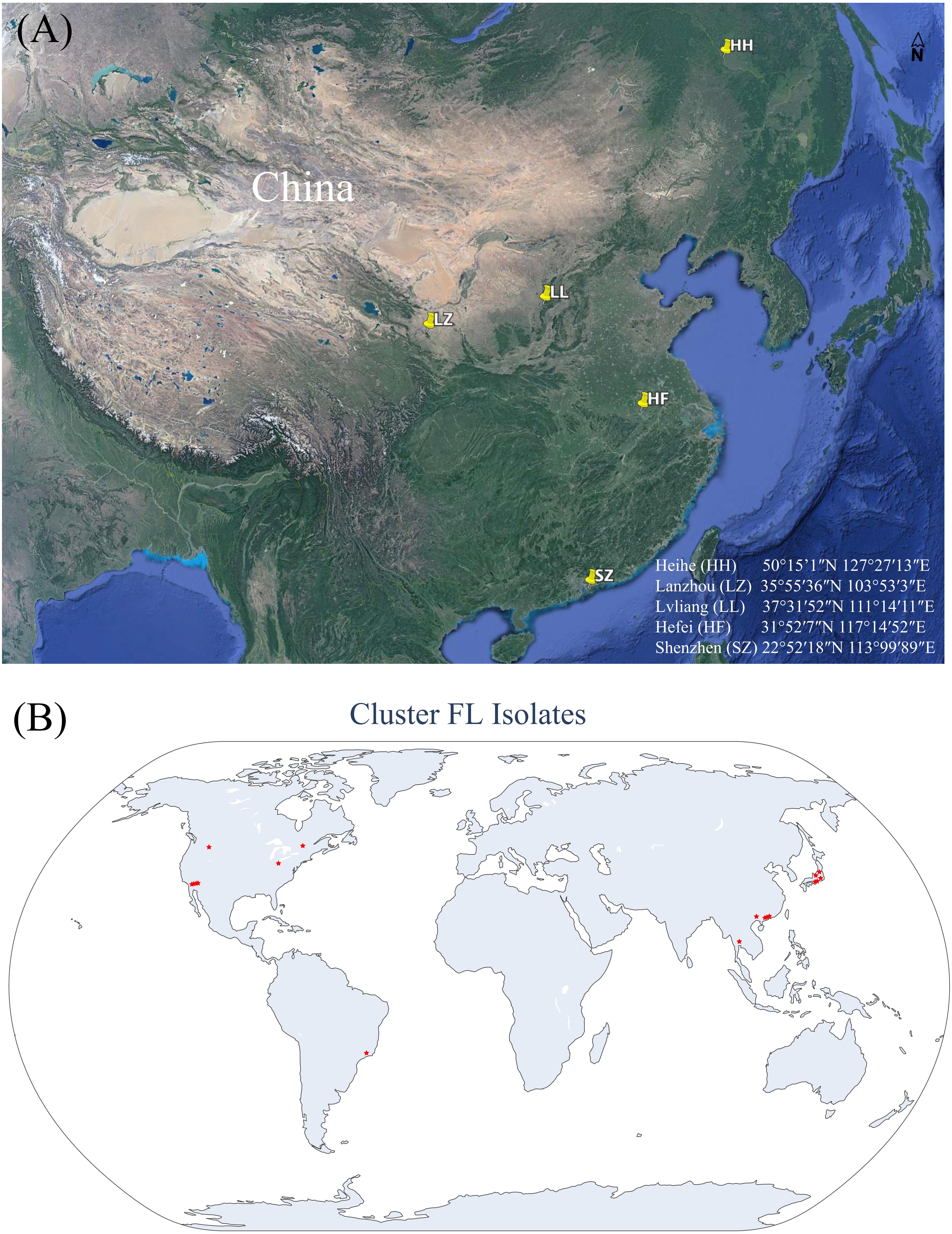


**Fig. S1**. Locations of isolation sites of *Bradyrhizobium*. (A) The locations of the five sampling sites in China where our 93 soil-dwelling *Bradyrhizobium* strains are isolated. The latitude and longitude of the sites are shown in the lower right corner. (B) The locations of the isolates belonging to Cluster FL in Fig. 2. Note that the isolation sites of a few strains are not known (see Data Set S2) and thus cannot be marked in the map.

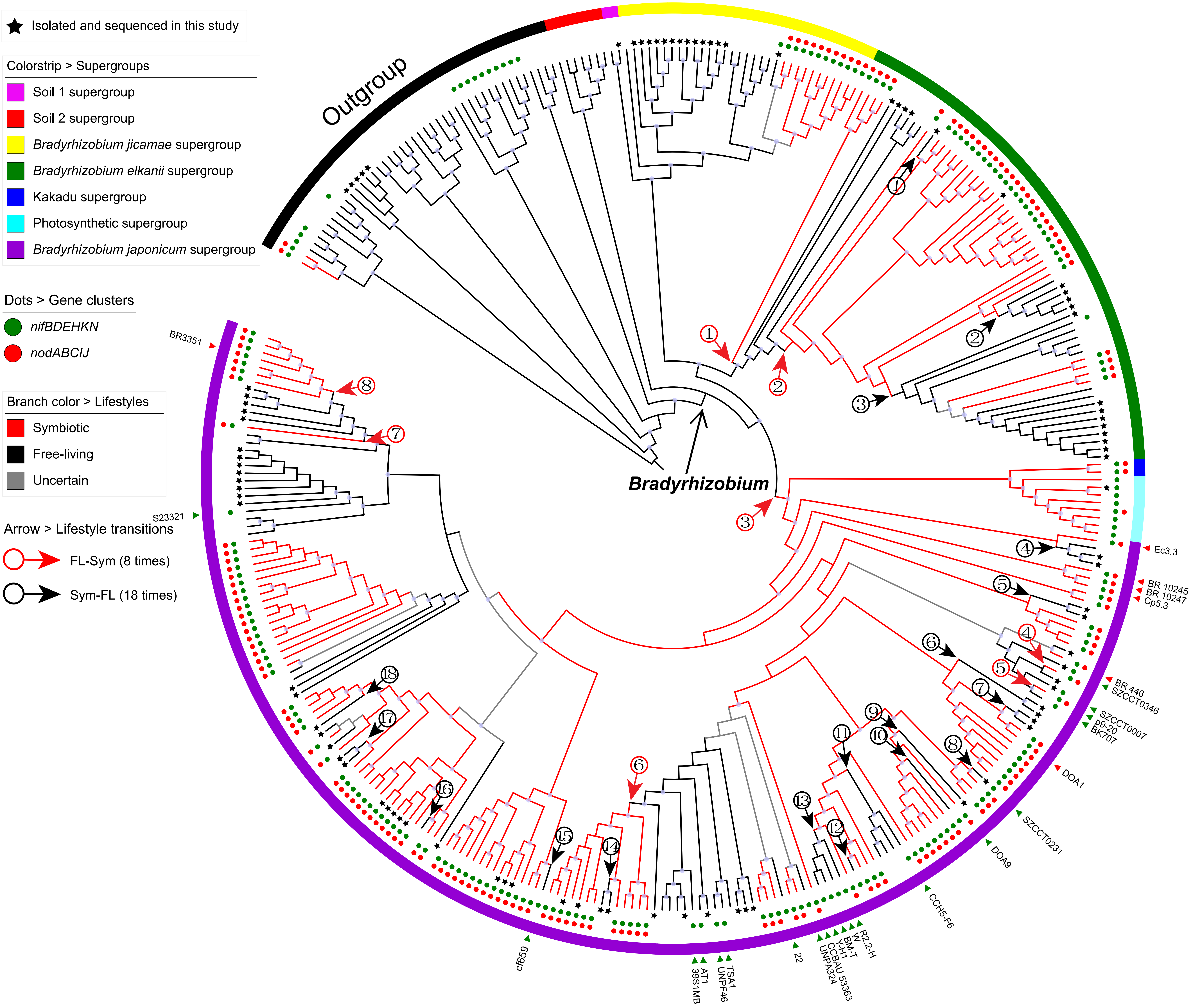


**Figure S2.** The maximum-likelihood phylogeny of *Bradyrhizobium* and inferred lifestyle evolutionary history. The meanings of the background colors, labels, and color strips are the same as shown in Fig. 1. We inferred 8 and 18 times FL-Sym and Sym-FL transitions respectively (noted with arrows).

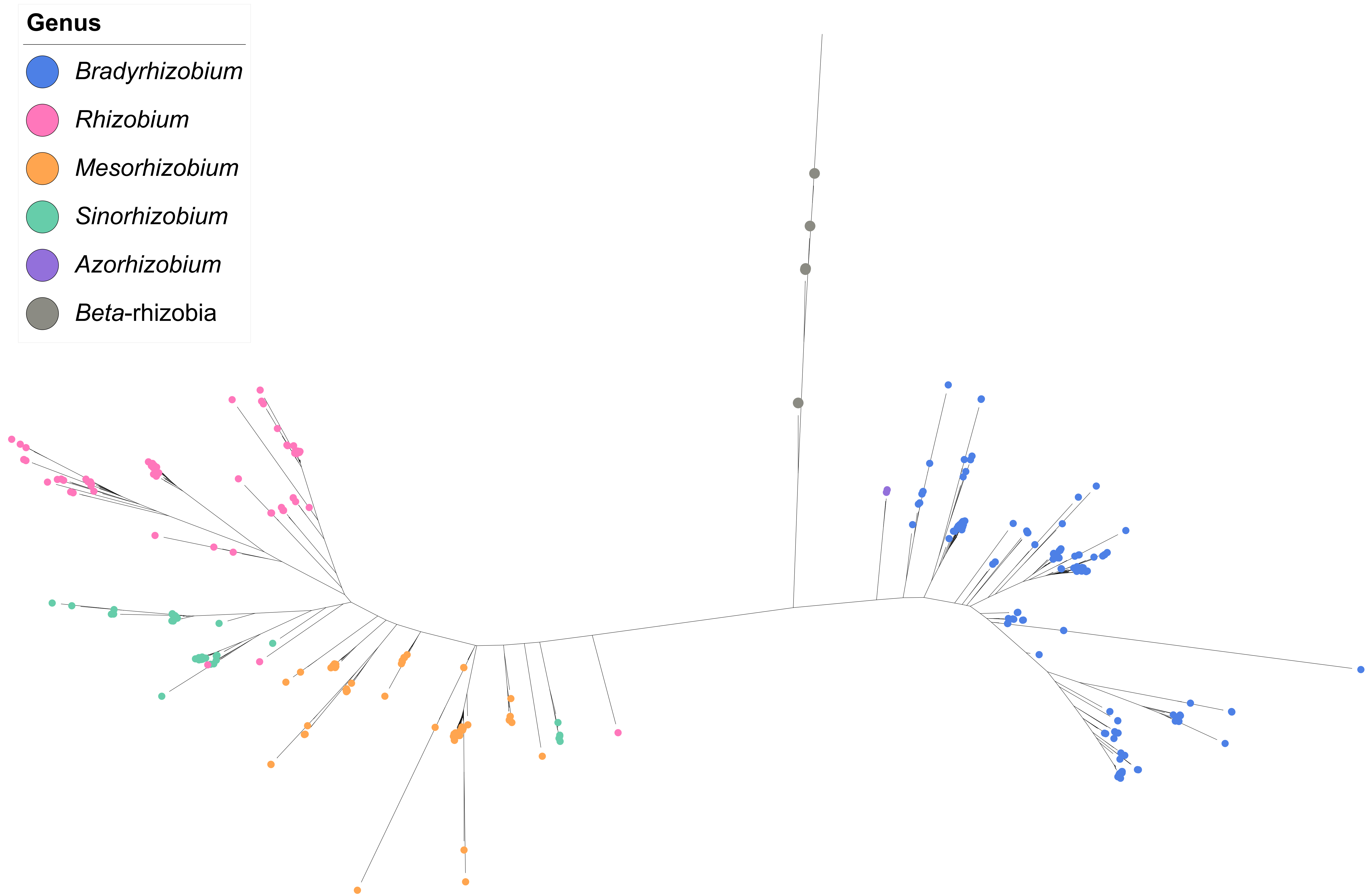


**Fig. S3**. Unrooted nif gene phylogeny of *Bradyrhizobium* and other rhizobia. Totally 436 strains are used for nif gene phylogeny constructing (244 in the outgroup). The phylogeny was constructed using the concatenated alignment of *nifABDEHKNVX* at the amino acid level. Nif genes from *Rhizobium*, *Sinorhizobium*, *Mesorhizobium*, *Azorhizobium*, and *Beta*-rhizobia are used as the outgroup. Different genera are denoted by circles with different colors.

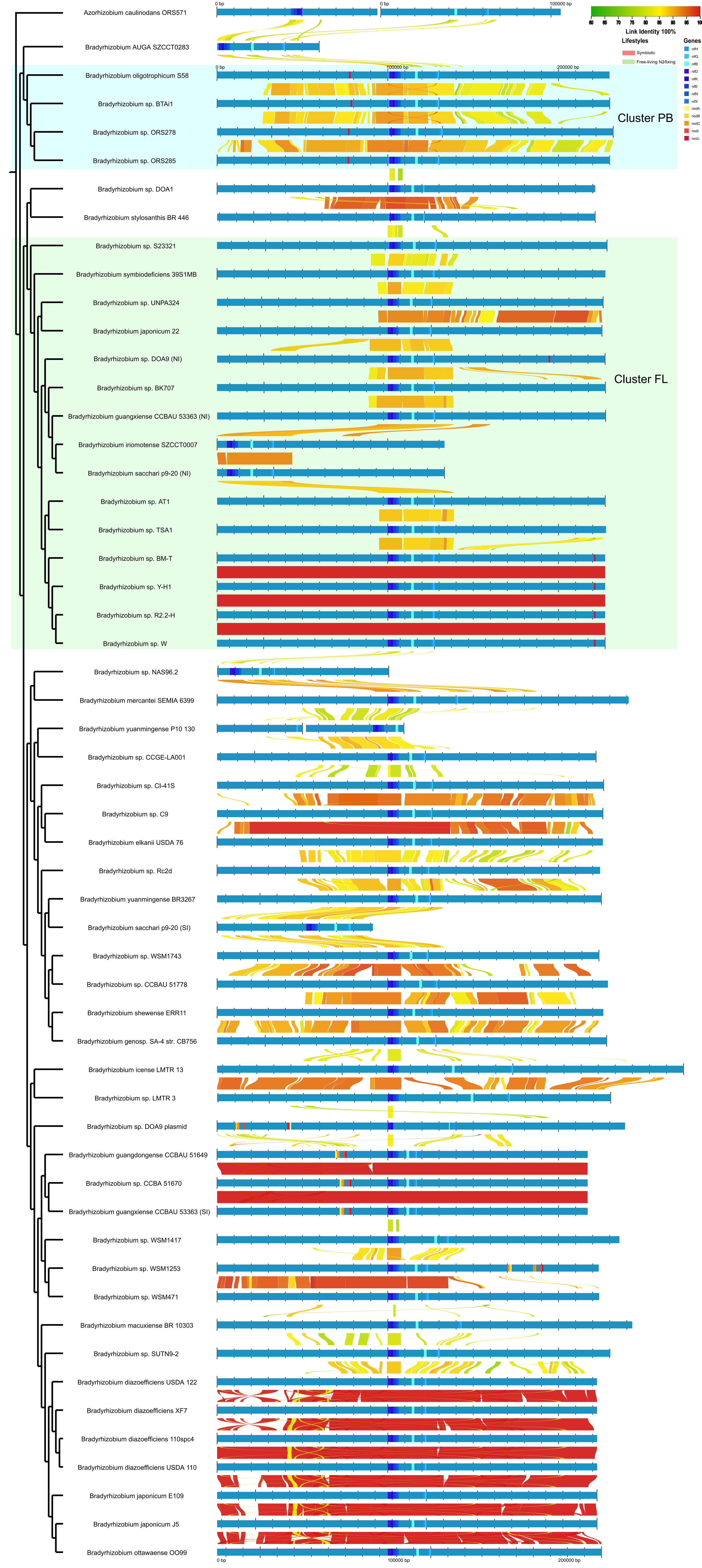


**Fig. S4**. Nucleotide sequence identities of DNA segments flanking nif genes. Blue bars represent the genome segment, and links between genomes indicate nucleotide identity. Genes are indicated by different colors in the blue bar. The background color under genome name indicates different lifestyles. The genomes were ordered according to their order in the nif gene tree. Genomes displayed here are i) those whose genome segment containing the nif genes is longer than 200 kb, and ii) those included in Fig. 3 (even if the nif-containing genome segment is shorter than 200 kb).

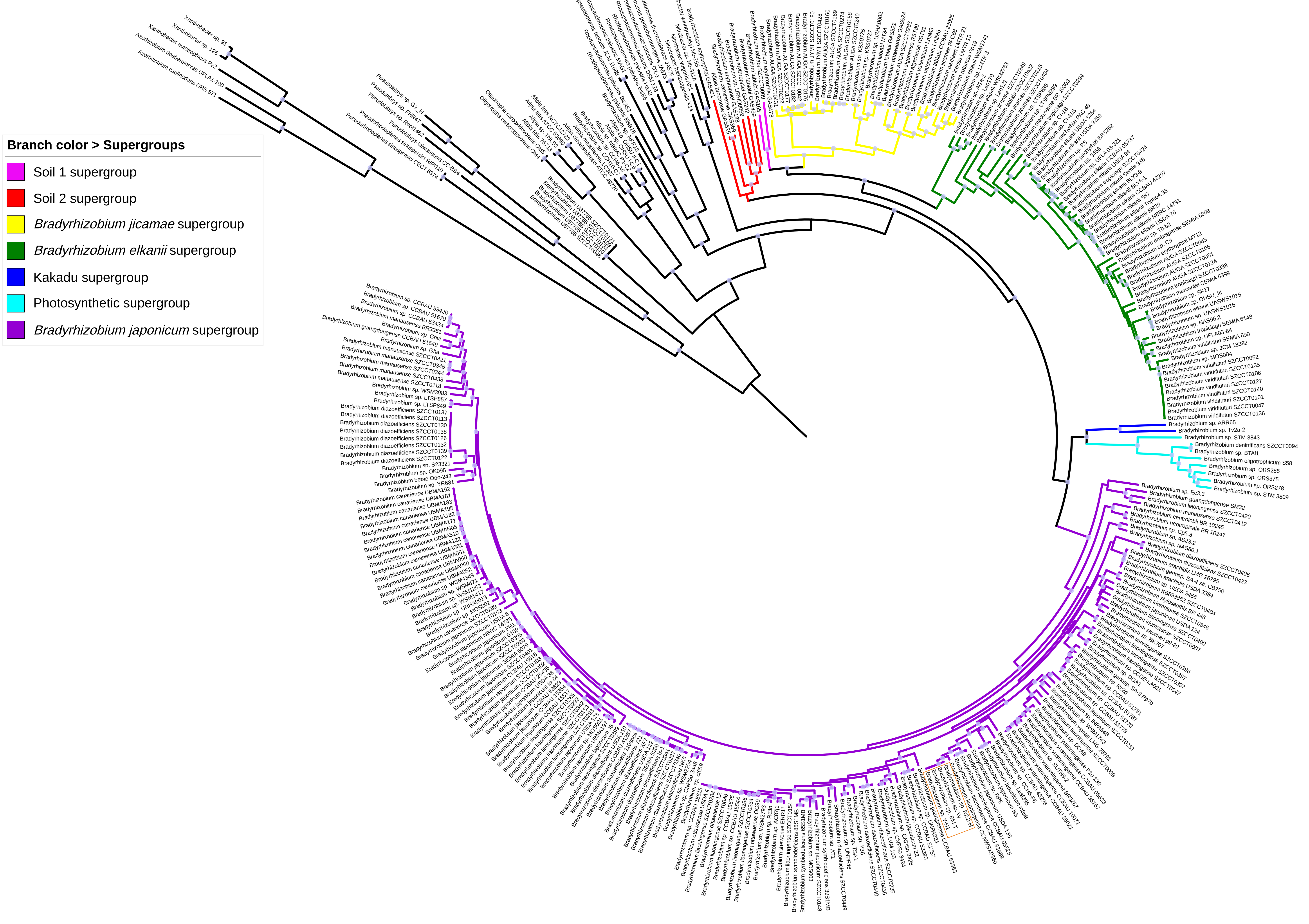


**Fig. S5**. The maximum-likelihood phylogenomic tree of *Bradyrhizobium*. The topology of the tree in this figure is the same as the one shown in Fig. 1 but with branch lengths representing substitution rates. The branch color indicates different supergroups. Purple circles on the nodes indicate ultrafast bootstrap values higher than or equal to 95% calculated by IQ-Tree.

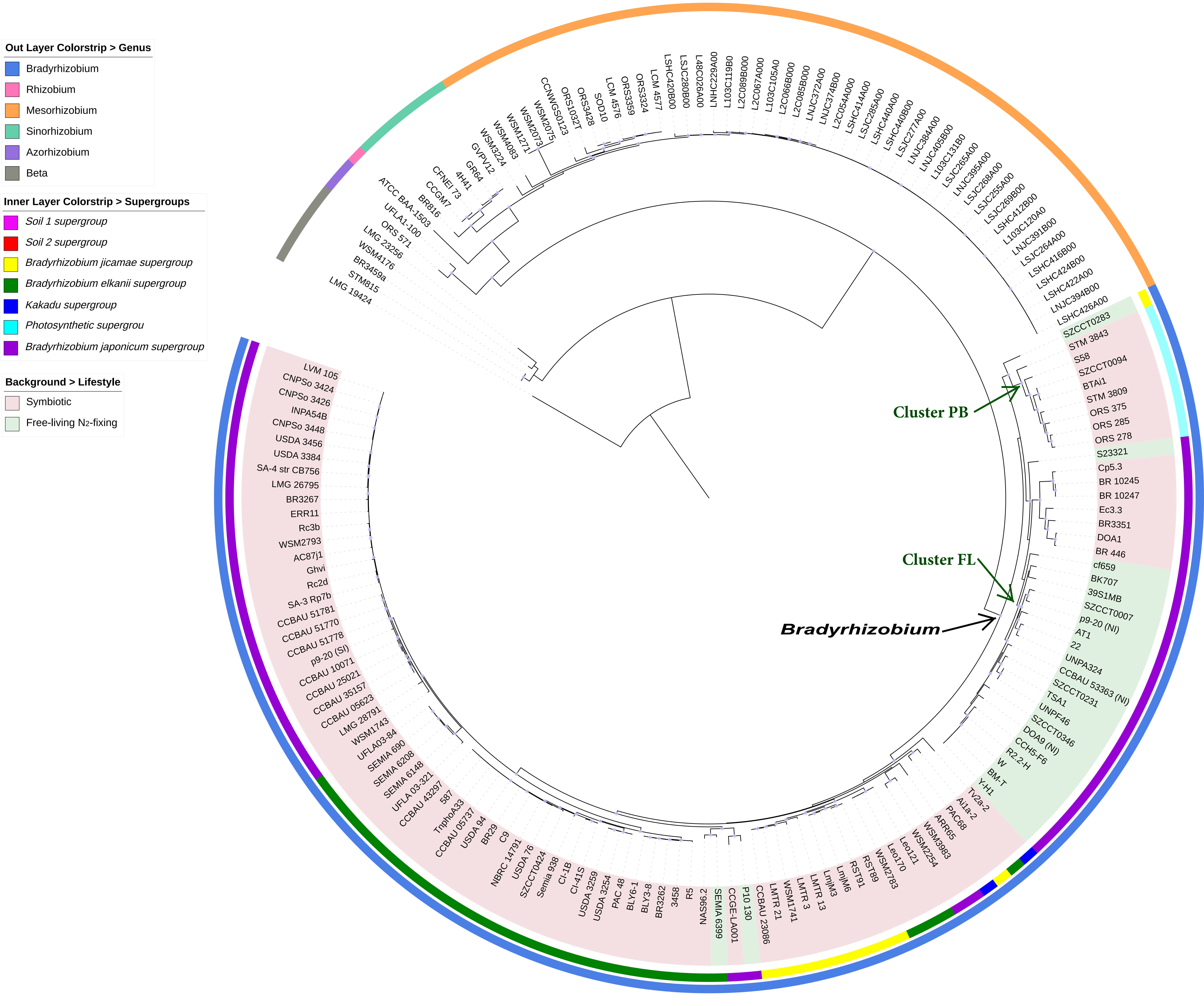


**Fig. S6**. *NifV* gene phylogeny of *Bradyrhizobium*. *NifV* from *Rhizobium*, *Sinorhizobium*, *Mesorhizobium*, *Azorhizobium* and *Beta*-rhizobia (rhizobia from *Betaproteobacteria*) are used as the outgroup. The color strips on the outer layer indicates different genes, color strips on the inner layer indicates different supergroups, and the background colors of the species name represent different lifestyles. The purple circles on the nodes indicate ultrafast bootstrap values higher than or equal to 95% calculated by IQ-Tree.

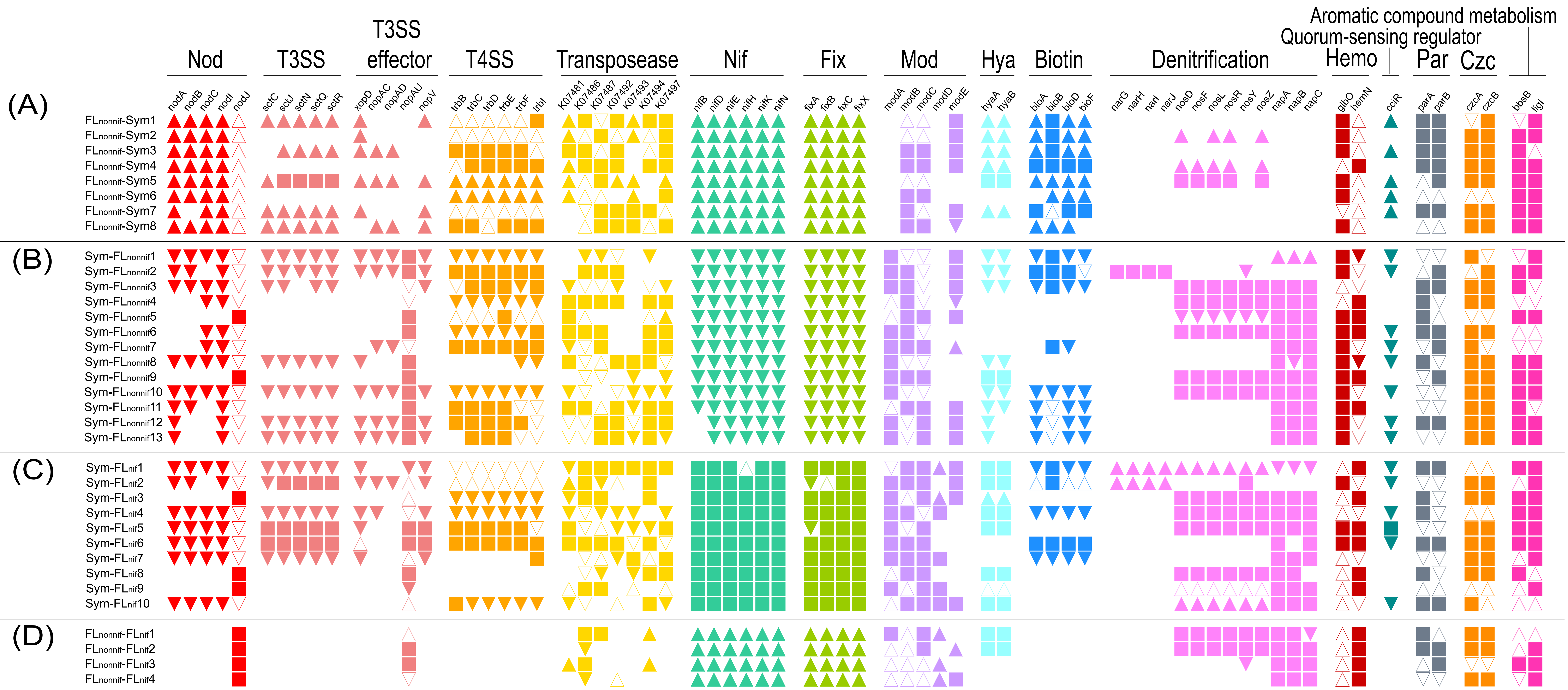


**Fig. S7**. Patterns of gene gains and losses during FL$_{nonnif}$-Sym (A), Sym-FL$_{nonnif}$ (B), Sym-FLnif (C) and FL$_{nonnif}$- FL$_{nif}$ (D) transitions. Events of gene gains and losses were inferred using BadiRate (see Materials and methods). A solid upward triangle indicates that the gene was absent in the ancestral node but gained during lifestyle transition, and an open upward triangle indicates that the gene was already present in ancestral nodes and its copy number increased during lifestyle transition. A solid downward triangle denotes the complete loss of the gene, while an open downward triangle denotes copy number contraction during lifestyle transition. A square indicates that the gene was present in the ancestral node and its copy number remained during lifestyle transition. The blanks indicate that corresponding genes were neither present in the ancestral nodes nor gained during lifestyle transition.

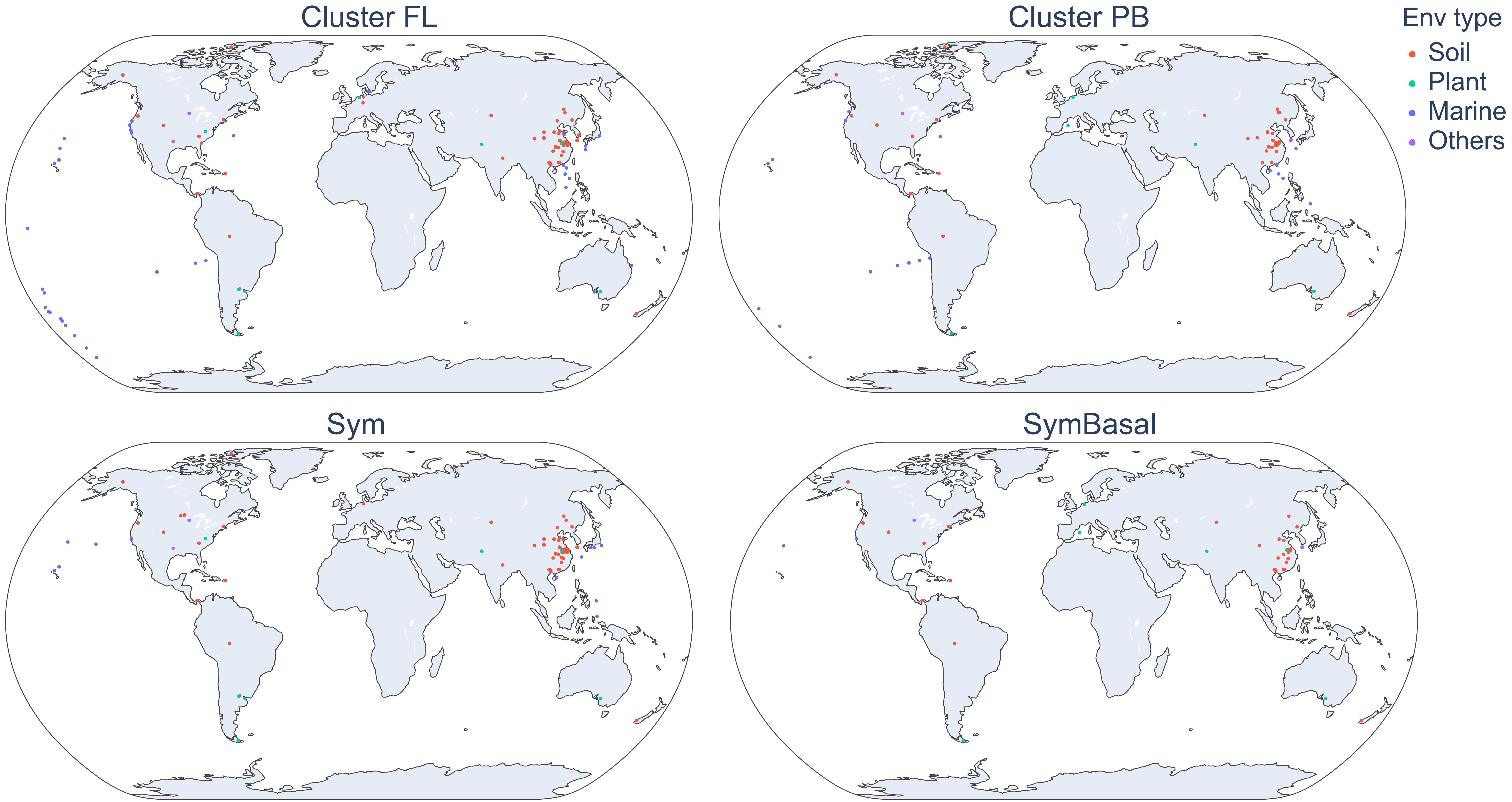


**Figure S8**. The geographical distribution of *nifH* based on the analysis of amplicon sequencing samples. Points marked on the map denote the sampling sites of *nifH* amplicon sequencing where the corresponding clusters, namely FL (free-living), PB (photosynthetic *Bradyrhizobium*), Sym (symbiotic), or Sym Basal (the basal group of symbiotic *nifH*) of *nifH* genes are detected (see Materials and methods).
